# Targeted Intracellular Degradation of SARS-CoV-2 RBD via Computationally-Optimized Peptide Fusions

**DOI:** 10.1101/2020.06.01.127829

**Authors:** Pranam Chatterjee, Manvitha Ponnapati, Joseph M. Jacobson

## Abstract

The COVID-19 pandemic, caused by the novel coronavirus SARS-CoV-2, has elicited a global health crisis of catastrophic proportions. With no approved cure or vaccine currently available, there is a critical need for effective antiviral strategies. In this study, we report a novel antiviral platform, through computational design of ACE2-derived peptides which both target the viral spike protein receptor binding domain (RBD) and recruit E3 ubiquitin ligases for subsequent intracellular degradation of SARS-CoV-2 in the proteasome. Our engineered peptide fusions demonstrate robust RBD degradation capabilities in human cells, thus prompting their further experimental characterization and therapeutic development.

## Introduction

SARS-CoV-2 has emerged as a highly pathogenic coronavirus that has now spread to over 200 countries, infecting nearly 6 million people worldwide and killing over 350,000 as of May 2020 (*1*). Economies have crashed, travel restrictions have been imposed, and public gatherings have been cancelled, all while a sizeable portion of the human population remains quarantined. Rapid transmission dynamics as well as a wide range of symptoms, from a simple dry cough to pneumonia and death, are common characteristics of coronavirus disease 2019 (COVID-19) (*2*). With no vaccine or cure readily available (*3*), there is a pressing need for robust and effective therapeutics targeting the virus.

Numerous antiviral strategies have been proposed to limit SARS-CoV-2 replication by preventing viral infection and synthesis (*4*). As SARS-CoV-2 is a positive-sense RNA virus, Abbott, et al. recently devised a CRISPR-Cas13d based strategy, termed PAC-MAN, to simultaneously degrade the positive-sense genome and viral mRNAs (*5*). While this method may serve as a potential prophylactic treatment, introducing foreign and relatively large components such as Cas13 enzymes into human cells *in vivo* presents various delivery and safety challenges (*6*).

The most rapid and acute method of protein degradation intracellularly is at the post-translational level. Specifically, E3 ubiquitin ligases can tag endogenous proteins for subsequent degradation in the proteasome (*7*). Thus, we hypothesize that by guiding E3 ubiquitin ligases to synthesized viral protein components, one can mediate depletion of SARS-CoV-2 in vivo, preventing further infection and replication.

In this study, we devise a targeted intracellular degradation strategy for SARS-CoV-2 by computationally designing peptides that bind to its spike (S) protein receptor binding domain (RBD) and recruit a human E3 ubiquitin ligase for subsequent proteasomal degradation. Our experimental results identify an optimal peptide variant that can mediate robust degradation of the RBD fused to a stable superfolder-green fluorescent protein (sfGFP) (*8*) in human cells, thus motivating further exploration of this strategy from a therapeutic perspective.

## Results

### Computationally-Optimized Peptides Targeting the SARS-CoV-2 RBD

Since the 2003 SARS epidemic, it has been widely known that the angiotensin-converting enzyme 2 (ACE2) receptor is critical for SARS-CoV entry into host cells (*9*). ACE2 is a monocarboxypeptidase, widely known for cleaving various peptides within the renin–angiotensin system (*10*). Functionally, there are two forms of ACE2. The full-length ACE2 contains a structural transmembrane domain, which anchors its extracellular domain to the plasma membrane. The extracellular domain has been demonstrated as a receptor for the S protein of SARS-CoV, and recently, for that of SARS-CoV-2 (*11*). The soluble form of ACE2 lacks the membrane anchor, thus preserving binding capacity, and circulates in small amounts in the blood (*12*).

Recently, it has been shown that soluble ACE2 (sACE2) can serve as a competitive interceptor of SARS-CoV-2 and other coronaviruses by preventing binding of the viral particle to the endogenous ACE2 transmembrane protein, and thus viral entry (*13*). sACE2, however, is capable of binding other biological molecules *in vivo*, most notably integrin receptors (*14*). Furthermore, it is critical that therapeutics targeting SARS-CoV-2 epitopes withstand the possibility of viral mutation, which may allow the virus to overcome the host adaptive immune response (*15*). We thus conducted in *silico* protein modeling to engineer minimal sACE2 peptides that not only maintain potent RBD binding, but also possess reduced off-target interaction with the integrin *α5β1* receptor, and exhibit cross-binding affinity toward the previous SARS-CoV spike protein, thus demonstrating tolerance to viral evolution (Figure 1A).

**Figure 1:**
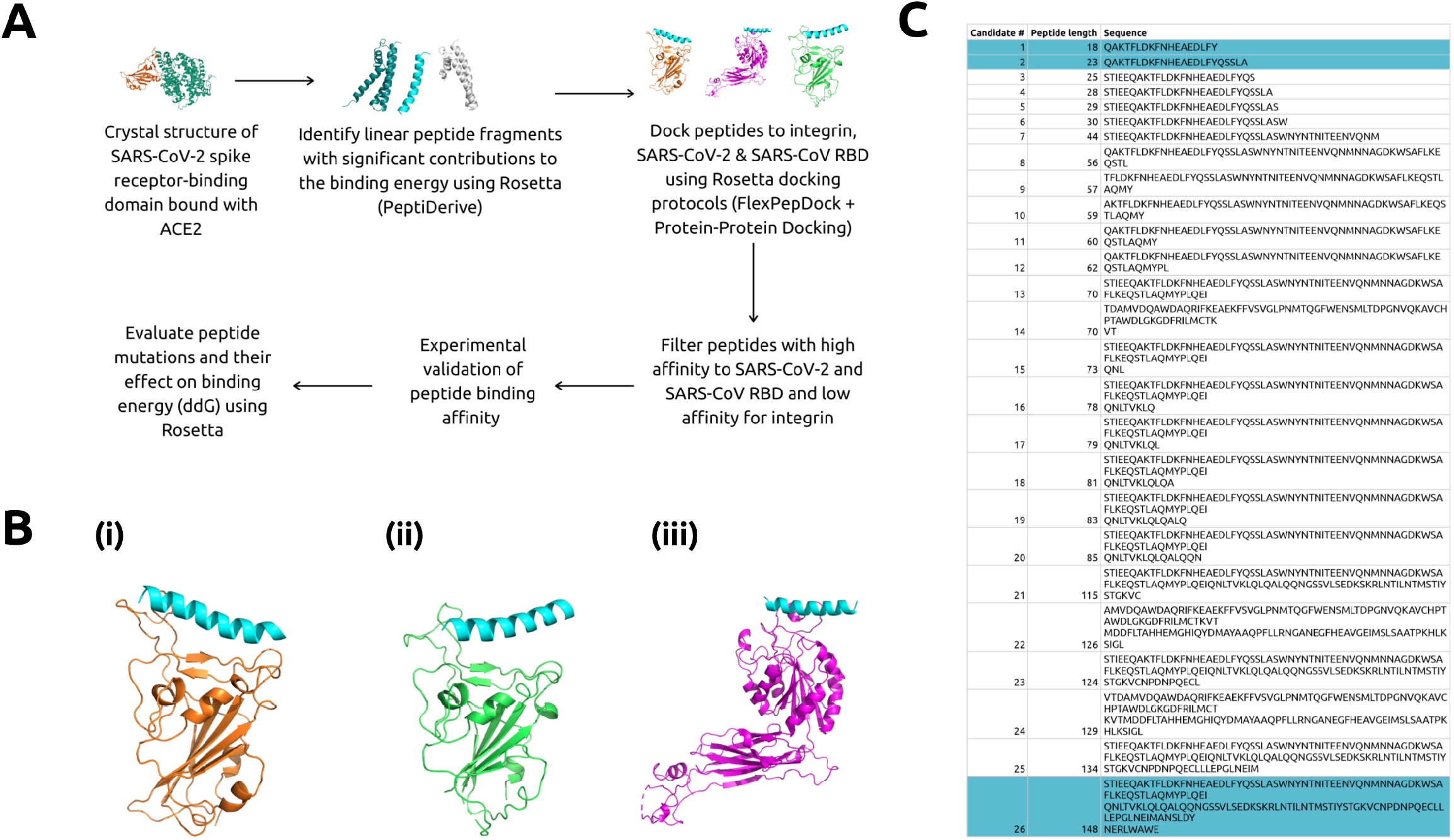
*In silico* Design of RBD-Targeting Peptides. A) Flow chart detailing computational pipeline to obtain optimized peptides. B) 23-mer peptide computationally docked to (i) SARS-CoV-2 RBD (*16*), (ii) SARS-CoV RBD (*20*), and (iii) integrin α5β1 receptor (*21*) in Rosetta and visualized using PyMol. The 23-mer peptide is shaded in blue. C) Candidate peptides selected after application of three filter docking steps. Peptides highlighted in blue indicate those chosen for experimental validation.

To do this, we retrieved a structure of the SARS-CoV-2 RBD bound to sACE2 from the Protein Data Bank (PDB 6M0J) (*16*). We first utilized the PeptiDerive protocol (*17*) in the Rosetta protein modeling software (*18*) to generate truncated linear sACE2 peptide segments between 10 to 150 amino acids with significant binding energy compared to that of the full SARS-CoV-2-sACE2-RBD interaction. To analyze the conformational entropy of the peptide segments in the binding pocket, we employed both the FlexPepDock and Protein-Protein protocols (*19*) to dock the peptides to the original RBD (Figure 1B.i). To ensure tolerance to potential mutations in the RBD, we docked peptides with optimal binding energies against the divergent 2003 SARS-CoV RBD bound with ACE2 (PDB 2AJF) (*20*) (Figure 1B.ii). Peptides that demonstrated highest binding energy for SARS-CoV and SARS-CoV-2 RBD were then docked against the α5β1 integrin ectodomain (PDB 3VI4) (*21*) to identify weak off-target binders (Figure 1B.iii). After applying these filters, 26 candidate peptides were selected from a total list of 188 initial peptides (Figure 1C).

### Targeted Degradation of RBD with TRIM21

TRIM21 is an E3 ubiquitin ligase that binds with high affinity to the Fc domain of antibodies and recruits the ubiquitin-proteasome system to degrade targeted proteins (*22*) Recently, the Trim-Away technique was developed for acute and rapid degradation of endogenous proteins, by co-expressing TRIM21 with an anti-target antibody (*23*). We thus hypothesized that by fusing the Fc domain to the C-terminus of candidate peptides and co-expressing TRIM21, we can mediate degradation of the RBD fused to a stable fluorescent marker, such as superfolder GFP (*8*) (RBD-sfGFP), in human HEK293T cells using a simple plasmid-based assay (Figure 2A and 2B). We chose the two most compact candidate peptides, an 18-mer and 23-mer derived from the ACE2 peptidase domain α1 helix, which is composed entirely of proteinogenic amino acids, as well as the candidate peptide computationally predicted to have highest binding affinity to the RBD (a 148-mer), for testing alongside sACE2 (Figure 1C). We also tested a recently-engineered 23-mer peptide from Zhang, et al., purporting to have strong RBD-binding capabilities (*24*). Five days post-transfection, we analyzed the degradation of the RBD-sfGFP complex by flow cytometry. After confirming negligible baseline depletion of GFP+ signal with and without exogenous TRIM21 expression, as well as no off-target degradation of sfGFP unbound to the RBD, we observed over 30% reduction of GFP+ cells treated with fulllength sACE2 fused to Fc and co-expressed with TRIM21, as compared to the RBD-sfGFP-only control. Of the tested peptides, only the 23-mer demonstrated comparable levels of degradation, with nearly 20% reduction in GFP+ cells (Figure 2C).

**Figure 2:**
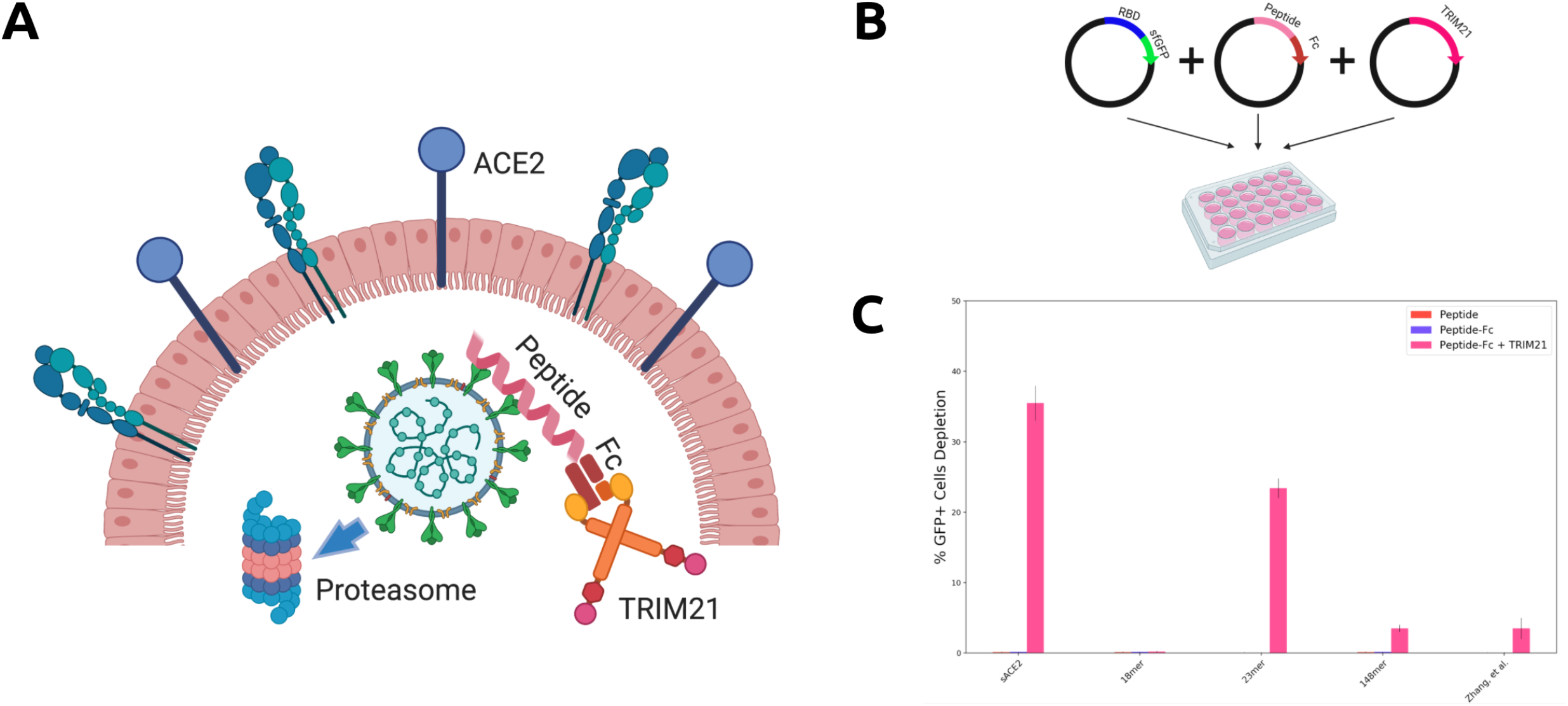
TRIM21-mediated Degradation of RBD via Peptide Targeting. A) Architecture and mechanism of TRIM2l-based degradation system. The Fc domain is fused to the C-terminus of RBD-targeting peptides. TRIM21 recognizes the Fc domain and tags SARS-CoV-2 for ubiquitin-mediated degradation in the proteasome. B) Three plasmid assay used to experimentally validate degradation architecture in human HEK293T cells. All CDS are inserted into the pcDNA3.1 backbone. C) Analysis of RBD-sfGFP degradation by flow cytometry, in the absence or presence of Fc (in *cis*), TRIM21 (in *trans*), or both. All samples were performed in independent transfection duplicates (n=2) and gated on GFP+ fluorescence. Mean percentage of GFP+ cell depletion was calculated in comparison to the RBD-sfGFP only control. Standard deviation was used to calculate error bars.

### Engineering of an Optimal Peptide-Based Degradation Architecture

Recently, deep mutational scans have been conducted on sACE2 to identify variants with higher binding affinity to the RBD of SARS-CoV-2 (*25*). Similarly, we conducted a complete single point mutational scan for all 23 positions in the peptide using the ddG-backrub script in Rosetta to identify mutants with improved binding affinity (*26*). For each mutation, 30,000 backrub trials were performed to sample conformational diversity. The top eight mutations predicted by this protocol were chosen for the experimental assay (Figure 3A.i), along with the top eight mutations predicted using an Rosetta energy function optimized for predicting the effect of mutations on protein-protein binding (*26*) (Figure 3A.ii), as well as the top eight mutational sites within the 23-mer sequence from deep mutational scans of sACE2 (*25*) (Figure 3A.iii). Our results in the subsequent TRIM21 assay identified A2N, derived from the original Rosetta energy function, as the optimal mutation in the 23-mer peptide, which achieved over 50% depletion of GFP+ cells, improving on both the sACE2 and 23-mer architecture as well as that of a previously optimized full-length mutant, sACE2v2.4 (Figure 3B).

**Figure 3:**
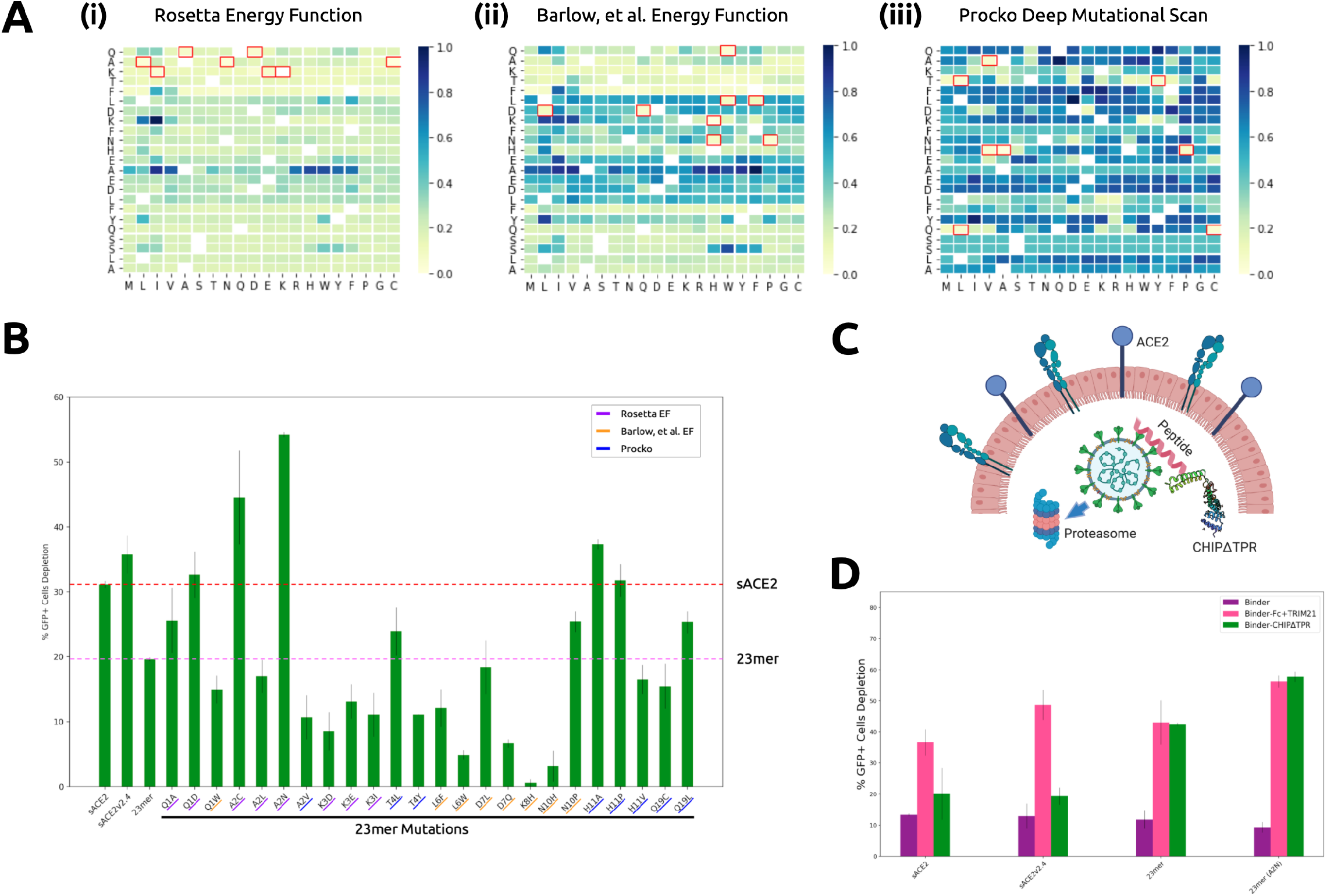
Engineering of Optimal RBD Degradation Architecture. A) (i) ddG of 23-mer mutations predicted by Rosetta with the default energy function, (ii) ddG of 23-mer mutations predicted by Rosetta with the energy function from Barlow et al. (*26*), (iii) log2fc enrichment scores of mutations of sACE2 within the 23-mer sequence experimentally determined by Procko (*25*). All binding affinity scores have been re-scaled to 0 (highest) to 1 (lowest) for visualization. Original amino acids are indicated on the y-axes. B) Analysis of RBD-sfGFP degradation by flow cytometry. All samples were performed in independent transfection duplicates (n=2) and gated on GFP+ fluorescence. All indicated samples were co-transfected with RBD-sfGFP and TRIM21 in *trans*, and mean percentage of GFP+ cell depletion was calculated in comparison to the RBD-sfGFP-only control. Standard deviation was used to calculate error bars. 23-mer mutations are underlined according to origin. C) Architecture and mechanism of CHIPΔTPR-based degradation system. CHIPΔTPR is fused to the C-terminus of RBD-targeting petpides. CHIPΔTPR can thus tag SARS-CoV-2 for ubiquitin-mediated degradation in the proteasome. D) Analysis of RBD-sfGFP degradation by flow cytometry in the presence of Fc (in *cis*) and TRIM21 (in *trans*) or CHIPΔTPR (in cis). All samples were performed in independent transfection duplicates (n=2) and gated on GFP+ fluorescence. Mean percentage of GFP+ cell depletion was calculated in comparison to the RBD-sfGFP only control. Standard deviation was used to calculate error bars.

Finally, numerous previous works have attempted to redirect E3 ubiquitin ligases by replacing their natural protein binding domains with those targeting specific proteins (*27–29*). In 2014, Portnoff, et al., reprogrammed the substrate specificity of a modular human E3 ubiquitin ligase called CHIP (carboxyl-terminus of Hsc70-interacting protein) by replacing its natural substrate-binding domain with designer binding proteins to generate optimized “ubiquibodies” or uAbs (*30*). To engineer a single construct that can mediate SARS-CoV-2 degradation without the need for trans expression of TRIM21, we fused the RBD-binding proteins to the CHIPΔTPR modified E3 ubiquitin ligase domain (Figure 3C). After co-transfection in HEK293T cells with the RBD-sfGFP complex, we observed that the 23-mer (A2N) mutant peptide maintained equivalent levels of degradation between the TRIM21 and CHIPΔTPR fusion architecture, and was more potent than that of sACE2, sACE2v2.4, and the original 23mer (Figure 3D).

## Discussion

In this study, we have computationally truncated and engineered the human ACE2 receptor sequence to potently bind to the SARS-CoV-2 RBD. We have further identified an optimized peptide variant that enables robust degradation of RBD-sfGFP complexes in human cells, both in trans and in cis with human E3 ubiquitin ligases. This mutant peptide not only maintains high binding affinity to the RBD, but is also predicted to bind to divergent RBD sequences in the case of future mutation as SARS-CoV-2 continues on its evolutionary trajectory. We also predict that the peptide will avoid binding to integrin α5β 1, thus preventing interference of critical cellular pathways.

While many further improvements and testing contexts are needed, there may be certain advantages to our platform as compared to the PAC-MAN strategy presented recently (*5*). First, both the peptide and E3 ubiquitin ligase components have been engineered from endogenous human proteins, unlike Cas13d, which is derived from *Ruminococcus flavefaciens* bacteria, thus potentially reducing the risk of immunogenicity. In terms of in vivo delivery as RNA or recombinant protein, Cas13d has an open reading frame (ORF) of nearly 1000 amino acids, not including the guide RNAs needed for interference. The entire peptide-CHIPΔTPR ORF consists of just over 200 amino acids, which can be readily synthesized as a peptide or be efficiently packaged for delivery in a lipid nanoparticle or adeno-associated virus (AAV). Similarly, the peptide-CHIPΔTPR complex can be utilized as a viral interceptor extracellularly, as it can competitively bind the RBD to prevent entry via ACE2. In the case that SARS-CoV-2 still infects the cell while bound to peptide-CHIPΔTPR, the virus would be immediately tagged for degradation in the proteasome.

In total, we envision that the strategy of utilizing a computationally-designed peptide binder linked to an E3 ubiquitin ligase can be explored not only for SARS-CoV-2, but also for other viruses that have known binding partners. Thus with further experimental characterization, the system presented here provides a potential new therapeutic platform in the fight against COVID-19 and future emergent viral threats.

## Materials and Methods

### Computational Peptide Design Pipeline

PDB Structure 6M0J containing the crystal structure of SARS-CoV-2 spike receptor-binding domain (RBD) bound with ACE2 was retrieved (*16*). The PeptiDerive protocol (*17*) in the Rosetta protein modeling software (*18*) was used to determine the linear peptide segments between 10 to 150 amino acids with significant binding energy compared to that of the whole SARS-CoV-2-ACE2-RBD interaction. To analyze the conformational entropy of the peptide segments in the binding pocket, a combination of FlexPepDock and Protein-Protein docking protocols (*19*) in Rosetta was used to dock the peptides to the original receptor binding domain. All peptides were placed in the binding pocket of SARS-CoV-2 RBD for local docking. Using FlexPepDock, 300 models were created for each peptide and the top 15 models were selected to calculate the score. The peptides with the best binding energies were docked against SARS-CoV RBD bound with ACE2 using PDB 2AJF (*20*) containing the crystal structure of SARS-CoV spike receptor-binding domain (RBD) bound with ACE2. Peptides that demonstrated highest binding energy for SARS-CoV and SARS-CoV-2 RBD were docked against the α5β 1 integrin ectodomain using the crystal structure provided by PDB 3VI4 (*21*).

The 23-mer peptide that showed high experimental binding affinity was selected for computational mutagenesis. A complete single point mutational scan was run for all 23 positions in the peptide using ddG-backrub script in Rosetta. For each mutation, 30,000 backrub trials were run to sample conformational diversity. The top 8 mutations predicted by this protocol were chosen for the experimental assay, along with the top 8 mutations predicted using an Rosetta energy function optimized for predicting the effect of muta-tions on protein-protein binding (*26*), as well as the top 8 mutational sites predicted by Procko on sACE2 (*25*).

### Generation of Plasmids

pcDNA3-SARS-CoV-2-S-RBD-sfGFP (Addgene Plasmid #141184) and pcDNA3-SARS-CoV-2-S-RBD-Fc (Addgene Plasmid #141183) were obtained as gifts from Erik Procko. hACE2 (Addgene Plasmid #1786) was obtained as a gift from Hyeryun Choe. Respective peptide DNA coding sequences (CDS) were amplified from hACE2 via PCR and inserted using HiFi DNA Assembly Master Mix (NEB) for Gibson Assembly into the pcDNA3-SARS-CoV-2-S-RBD-Fc backbone linearized by digestion with NheI and BamHI. pLVX puro TRIM21-GFP (Addgene Plasmid #1786) was obtained as a gift from Gaudenz Danuser. The TRIM21 CDS was amplified with overhangs for Gibson Assembly-mediated insertion into the pcDNA3-SARS-CoV-2-S-RBD-Fc backbone linearized by digestion with NheI and XhoI. pcDNA3-R4-uAb (Plasmid #101800) was obtained as a gift from Matthew DeLisa. Candidate sACE2 sequences were amplified from hACE2 with overhangs for Gibson Assembly-mediated insertion into linearized pcDNA3-R4-uAb digested with HindIII and EcoRI. Single amino acid substitutions were introduced utilizing the KLD Enzyme Mix (NEB) following PCR amplification with mutagenic primers (Genewiz). Assembled constructs were transformed into 50 *μ*L NEB Turbo Competent E. coli cells, and plated onto LB agar supplemented with the appropriate antibiotic for subsequent sequence verification of colonies and plasmid purification.

### Cell Culture and FACS Analysis

HEK293T cells were maintained in Dulbecco’s Modified Eagle’s Medium (DMEM) supplemented with 100 units/ml penicillin, 100 mg/ml streptomycin, and 10% fetal bovine serum (FBS). RBD-sfGFP (333 ng), peptide (333 ng), and E3 ubiquitin ligase (333 ng) plasmids were transfected into cells as duplicates (2 x 10^5^/well in a 24-well plate) with Lipofectamine 3000 (Invitrogen) in Opti-MEM (Gibco). For transfection conditions with fewer required plasmids, we co-transfected cells with the necessary amounts of empty pcDNA3.1 plasmid to obtain 999 ng of total DNA. After 5 days post-transfection, cells were harvested and analyzed on an Attune^®^ NxT Flow Cytometer (Thermo Fisher) for GFP fluorescence (488-nm laser excitation, 530/30 filter for detection). Cells expressing GFP were gated, and percent GFP+ depletion to the RBD-sfGFP only control were calculated. All samples were performed in independent transfection duplicates (n=2), and percentage depletion values were averaged. Standard deviation was used to calculate error bars.

## Acknowledgments

We thank Dr. Neil Gershenfeld and Dr. Shuguang Zhang for shared lab equipment. We further thank Eyal Perry for computational assistance, and James Weis and Nest.Bio Labs for access to flow cytometry.

## Author Contributions

P.C. conceived, designed, and supervised the study. P.C. designed and built constructs, carried out experiments, and conducted data analyses. M.P. implemented computational design pipelines and performed in *silico* docking protocols. P.C. and M.P. wrote the paper. J.M.J. reviewed the paper and provided critical insight and ideas.

## Declarations

This work was supported by the consortia of sponsors of the MIT Media Lab and the MIT Center for Bits and Atoms. P.C. and J.M.J. are listed inventors for provisional patent application entitled “Minimal Peptide Fusions for Targeted Intracellular Protein Degradation.”

## Data and Materials Availability

All data needed to evaluate the conclusions in the paper are present in the paper. Raw computational and experimental data files can be found at: shorturl.at/bipH3.

## References

1. Dong, E., Du, H. & Gardner, L. An interactive web-based dashboard to track COVID-19 in real time. The Lancet Infectious Diseases 20, 533–534 (2020).

2. Wu, J. T. et al. Estimating clinical severity of COVID-19 from the transmission dynamics in wuhan, china. Nature Medicine 26, 506–510 (2020).

3. Lurie, N., Saville, M., Hatchett, R. & Halton, J. Developing covid-19 vaccines at pandemic speed. New England Journal of Medicine 382, 1969–1973 (2020).

4. Senanayake, S. L. Drug repurposing strategies for COVID-19. Future Drug Discovery 2 (2020).

5. Abbott, T. R. et al. Development of CRISPR as an antiviral strategy to combat SARS-CoV-2 and influenza. Cell 181, 865–876.e12 (2020).

6. Lino, C. A., Harper, J. C., Carney, J. P. & Timlin, J. A. Delivering CRISPR: a review of the challenges and approaches. Drug Delivery 25, 1234–1257 (2018).

7. Ardley, H. C. & Robinson, P. A. E3 ubiquitin ligases. Essays in Biochemistry 41, 15–30 (2005).

8. Pédelacq, J.-D., Cabantous, S., Tran, T., Terwilliger, T. C. & Waldo, G. S. Engineering and characterization of a superfolder green fluorescent protein. Nature Biotechnology 24, 79–88 (2005).

9. Du, L. et al. The spike protein of SARS-CoV — a target for vaccine and therapeutic development. Nature Reviews Microbiology 7, 226–236 (2009).

10. Tipnis, S. R. et al. A human homolog of angiotensin-converting enzyme. Journal of Biological Chemistry 275, 33238–33243 (2000).

11. Zhou, P. et al. A pneumonia outbreak associated with a new coronavirus of probable bat origin. Nature 579, 270–273 (2020).

12. Wysocki, J. et al. Targeting the degradation of angiotensin II with recombinant angiotensin-converting enzyme 2. Hypertension 55, 90–98 (2010).

13. Monteil, V. et al. Inhibition of SARS-CoV-2 infections in engineered human tissues using clinical-grade soluble human ACE2. Cell 181, 905–913.e7 (2020).

14. Clarke, N. E., Fisher, M. J., Porter, K. E., Lambert, D. W. & Turner, A. J. Angiotensin converting enzyme (ACE) and ACE2 bind integrins and ACE2 regulates integrin signalling. PLoS ONE 7, e34747 (2012).

15. Andersen, K. G., Rambaut, A., Lipkin, W. I., Holmes, E. C. & Garry, R. F. The proximal origin of SARS-CoV-2. Nature Medicine 26, 450–452 (2020).

16. Lan, J. et al. Structure of the SARS-CoV-2 spike receptor-binding domain bound to the ACE2 receptor. Nature 581, 215–220 (2020).

17. Sedan, Y., Marcu, O., Lyskov, S. & Schueler-Furman, O. Peptiderive server: derive peptide inhibitors from protein–protein interactions. Nucleic Acids Research 44, W536–W541 (2016).

18. Rohl, C. A., Strauss, C. E., Misura, K. M. & Baker, D. Protein structure prediction using rosetta. Methods in Enzymology 66–93 (2004).

19. London, N., Raveh, B., Cohen, E., Fathi, G. & Schueler-Furman, O. Rosetta FlexPep-Dock web server—high resolution modeling of peptide–protein interactions. Nucleic Acids Research 39, W249–W253 (2011).

20. Li, F. Structure of SARS coronavirus spike receptor-binding domain complexed with receptor. Science 309, 1864–1868 (2005).

21. Nagae, M. et al. Crystal structure of 51 integrin ectodomain: Atomic details of the fibronectin receptor. The Journal of Cell Biology 197, 131–140 (2012).

22. Foss, S. et al. TRIM21—from intracellular immunity to therapy. Frontiers in Immunology 10 (2019).

23. Clift, D. et al. A method for the acute and rapid degradation of endogenous proteins. Cell 171, 1692–1706.e18 (2017).

24. Zhang, G., Pomplun, S., Loftis, A. R., Loas, A. & Pentelute, B. L. The first-in-class peptide binder to the SARS-CoV-2 spike protein. bioRxiv (2020).

25. Procko, E. The sequence of human ACE2 is suboptimal for binding the s spike protein of SARS coronavirus 2. bioRxiv (2020).

26. Barlow, K. A. et al. Flex ddG: Rosetta ensemble-based estimation of changes in protein-protein binding affinity upon mutation. The Journal of Physical Chemistry B 122, 5389–5399 (2018).

27. Gosink, M. M. & Vierstra, R. D. Redirecting the specificity of ubiquitination by modifying ubiquitin-conjugating enzymes. Proceedings of the National Academy of Sciences 92, 9117–9121 (1995).

28. Zhou, P., Bogacki, R., McReynolds, L. & Howley, P. M. Harnessing the ubiquitination machinery to target the degradation of specific cellular proteins. Molecular Cell 6, 751–756 (2000).

29. Su, Y., Ishikawa, S., Kojima, M. & Liu, B. Eradication of pathogenic-catenin by skp1/cullin/f box ubiquitination machinery. Proceedings of the National Academy of Sciences 100, 12729–12734 (2003).

30. Portnoff, A. D., Stephens, E. A., Varner, J. D. & DeLisa, M. P. Ubiquibodies, synthetic e3 ubiquitin ligases endowed with unnatural substrate specificity for targeted protein silencing. Journal of Biological Chemistry 289, 7844–7855 (2014).

